# Discovery of high-specificity DNA aptamers for progesterone using a high-throughput array platform

**DOI:** 10.1101/2025.09.13.675901

**Authors:** Hajime Fujita, Leighton Wan, Yasser Gidi, Linus A. Hein, Michael Eisenstein, Hyongsok Tom Soh

## Abstract

Aptamer-based biosensors offer several advantages for detecting small molecules, including chemical stability and compatibility with diverse sensing formats. However, developing highly specific DNA aptamers that can distinguish between structurally similar small-molecule analytes remains a major challenge. Steroid hormones share a common four-ring scaffold, and even small modifications in functional groups can lead to distinct biological activities. Although DNA aptamers targeting the female reproductive hormone progesterone have been previously reported, many exhibit substantial cross-reactivity with other steroid hormones. In this work, we report the discovery of DNA aptamers with high affinity and specificity for progesterone using our aptamer array platform, which can characterize millions of aptamer candidates in a single, automated experiment. This platform allows us to profile aptamer libraries from earlier rounds of systematic evolution of ligands by exponential enrichment (SELEX), while preserving sequence diversity. Using this strategy, we identified multiple aptamers with nanomolar affinity for progesterone and minimal cross-reactivity to structurally related steroids. These high-specificity aptamers provide a strong foundation for the development of biosensors applicable to both clinical diagnostics and biological research.

## Introduction

DNA aptamers are nucleic acid-based affinity reagents that bind molecular targets with high affinity and specificity^1,2^. Their chemical stability and ease of engineering for binding-induced signal readout make them useful tools for biosensing and diagnostic applications^3–7^. However, it remains challenging to develop aptamers that can bind tightly to and distinguish between chemically similar small-molecule analytes^8,9^. For small-molecule targets, conventional aptamer selection via systematic evolution of ligands by exponential enrichment (SELEX) typically requires 15 or more rounds, with careful adjustment of target and counter-target concentrations throughout the process^10,11^. While early-round SELEX pools may contain high-affinity candidates, they are rarely evaluated functionally, as conventional workflows typically focus on sequencing and characterization only after extensive enrichment^12,13^.

To overcome these limitations, our lab has previously developed a high-throughput aptamer array platform that enables parallel binding measurements of millions of sequences in a single run^2,14,15^. This platform allows us to sequentially flow in multiple related compounds and measure their effects in a single run, enabling simultaneous profiling of affinity and specificity. By applying this approach to early-stage SELEX libraries (*e*.*g*., rounds 5–10), we can evaluate functional performance before sequence convergence occurs, allowing for earlier and more informed aptamer selection. We previously validated this strategy by conducting only seven rounds of SELEX to identify DNA aptamers for tryptophan metabolites. Remarkably, the selected aptamers could discriminate between targets differing by just a single hydroxyl group^13,14^. Encouraged by these results, we have now applied the same strategy to steroid hormones—a more clinically relevant yet chemically challenging target class for aptamer discovery.

Here, we report the discovery of high-affinity, high-specificity DNA aptamers for progesterone using this array-based screening platform. Accurate detection of progesterone is essential for fertility tracking and reproductive health management, as its levels fluctuate substantially over short time intervals, ranging from low to high nanomolar concentrations within hours in ovulatory women^16,17^. However, prior DNA aptamers for progesterone have exhibited substantial cross-reactivity with analogs such as estradiol^18,19^. Furthermore, most prior screens did not assess cross-reactivity with cortisol, a structurally similar and physiologically abundant hormone. By incorporating such non-target interferents directly into our multiplexed screening runs, we were able to identify aptamers with strong responses to progesterone and minimal responses to analogs. Two top candidates, PRO-01 and PRO-02, exhibited dissociation constants (*K*_*D*_) of 28 nM and 36 nM, and negligible binding to non-target steroids. This work highlights the utility of our platform for discovering aptamers with high affinity and specificity, even for classes of targets that are chemically subtle yet biologically complex. The high specificity of the selected aptamers supports the development of biosensors for clinical and research applications, including accurate monitoring of fertility and reproductive health.

## Results and Discussion

### Screening approach for isolating high-affinity, target-specific aptamers

We sought to generate a DNA aptamer with high affinity and specificity for the steroid hormone progesterone using our previously reported high-throughput aptamer screening pipeline that combines capture-SELEX with a custom aptamer array platform^2,14,15^. In capture-SELEX, a DNA library consisting of random sequence regions flanked by constant primer regions is hybridized to complementary biotinylated capture strands and immobilized on a streptavidin-coated resin column (**Fig. 1a**). After applying a solution containing progesterone to the resin-immobilized aptamers, sequences that bind to the target undergo a conformational change that results in dissociation from the capture strand and elution into the solution. This structure-switching mechanism ensures that only sequences with high affinity for the target are recovered and amplified for subsequent rounds of selection.

**Figure 1.**
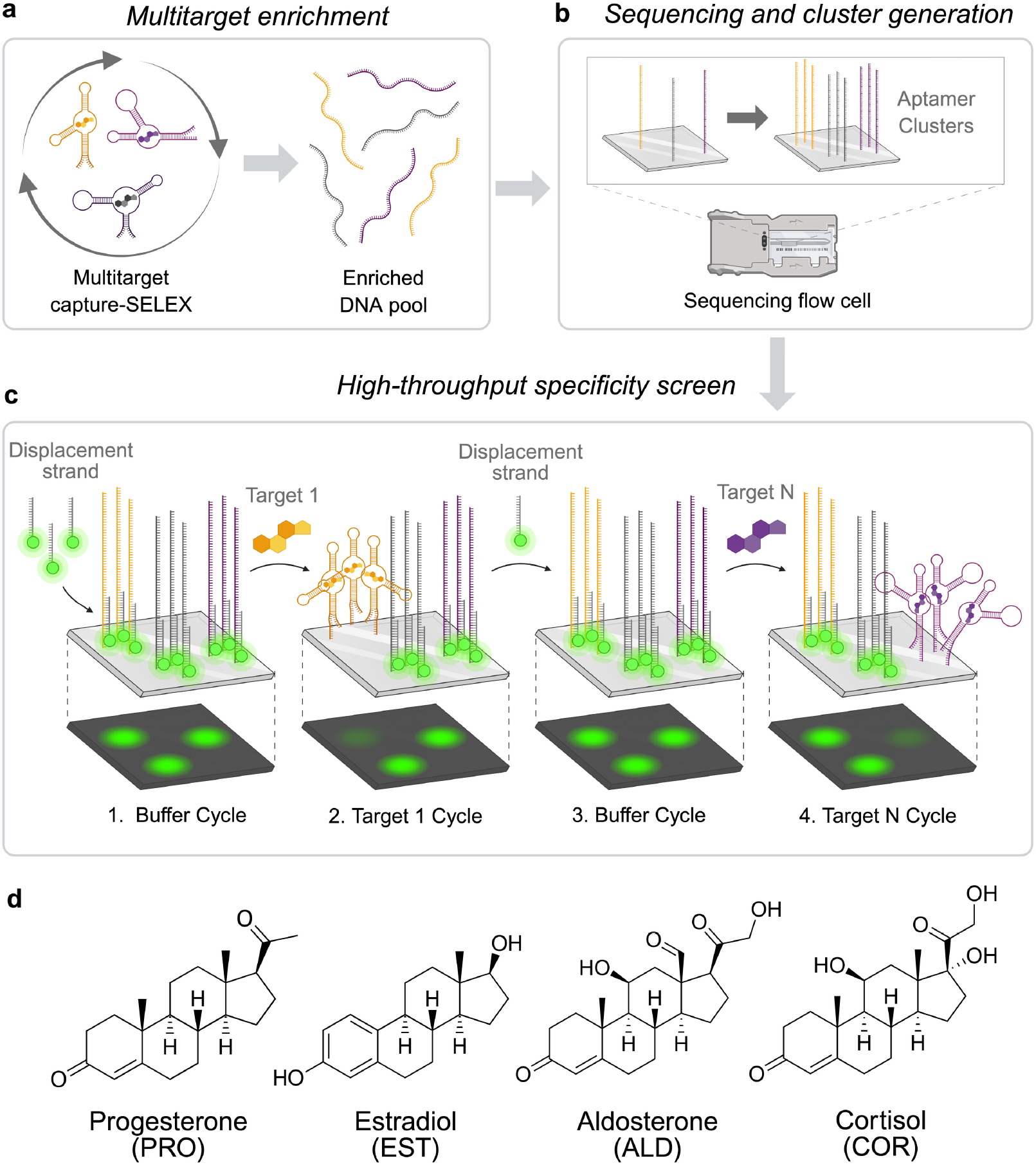
Overview of our high-throughput screening pipeline for aptamers with high affinity and specificity for progesterone. (**a**) Capture-SELEX workflow for isolating aptamer candidates with target-induced structure switching from a randomized DNA library. (**b**) Cluster generation and sequencing on Illumina MiSeq. (**c**) Integrated analysis of binding affinity and specificity using alternating buffer and target incubation cycles. Fluorescence from a fluorophore-labeled displacement strand is measured before and after target incubation. (**d**) Chemical structures of the steroid analytes used in screening: progesterone (PRO), estradiol (EST), aldosterone (ALD) and cortisol (COR).

Following enrichment, we screened aptamer candidates on our high-throughput aptamer array platform designed to evaluate binding performance at scale. This platform enables the simultaneous measurement of both affinity and specificity across millions of sequences by combining fluorescence-based binding assays with high-throughput sequencing. As shown in **Figure 1b**, aptamers are immobilized on the surface of an Illumina MiSeq flow cell and then sequenced to establish the identity of every sequence at each coordinate on the flow cell. Subsequent imaging cycles with and without target analytes enable fluorescence-based detection of ligand binding at each location (**Fig. 1c**).

We first apply progesterone to the flow cell and measure the loss of fluorescence due to displacement of a fluorophore-labeled DNA strand. This readout reflects aptamer conformational switching upon ligand binding. By repeating this process with structurally similar non-target analytes, we directly compare affinity and specificity across the library (**Fig. 1d**). This enables the rapid identification of aptamers that bind strongly and selectively to progesterone while discriminating against analogs such as estradiol, aldosterone, and cortisol.

### Screening progesterone aptamers using a high-throughput aptamer array platform

We first enriched the aptamer pool using capture-SELEX and then assessed the binding performance of enriched aptamers on our high-throughput array. The starting DNA library for capture-SELEX consisted of a central 30-nucleotide (nt) randomized region flanked by 15-nt constant primer regions (**Fig. 1a, Table S1; SI Section i.b**). During PCR amplification of the eluted pool, a Cy3-labeled forward primer was used, generating fluorescently labeled DNA products (**SI Section i.d**). This fluorescent labeling enabled us to monitor aptamer enrichment across rounds by measuring the fluorescence of eluted fractions. We observed a sharp increase in signal between rounds 5 and 7, indicating enrichment of aptamers that undergo target-induced release from the capture strand.

Following enrichment, we employed a modified Illumina MiSeq sequencer to profile sequence-affinity relationships within the aptamer pool^2,14^. The pre-enriched DNA sequences from round 5 of the capture-SELEX were immobilized onto the sequencer flow cell surface, where bridge amplification generated spatially localized clusters of identical sequences (**Fig. 1b; SI Section i.e**,**f**). We observed 1.7 million aptamer clusters on the flow cell, enabling parallel measurement of their binding interactions in a single experiment. Each aptamer cluster was then hybridized with Cy3-labeled DNA displacement strands that were identical to the capture strand (15 nt) used in SELEX (**Fig. 1c**). We then applied 50 µM progesterone to the flow cell, and target-binding aptamers underwent a conformational change that led to release of the Cy3-labeled displacement strands, yielding a measurable reduction in fluorescent signal. We normalized these changes to buffer-only controls and converted them to Z-scores for each aptamer cluster (**Fig. 2a– d**). Z-scores were calculated by first subtracting the mean percent change of all clusters within a given cycle from each individual cluster’s percent change, and then dividing the result by the standard deviation of percent changes for that cycle (**SI Section i.g**). To assess specificity, we extended this approach by performing additional imaging cycles using structurally-similar steroids such as estradiol (10 µM), cortisol (100 µM), and aldosterone (100 µM). For each analyte, we calculated Z-scores based on the fluorescence difference between target and buffer conditions.

**Figure 2.**
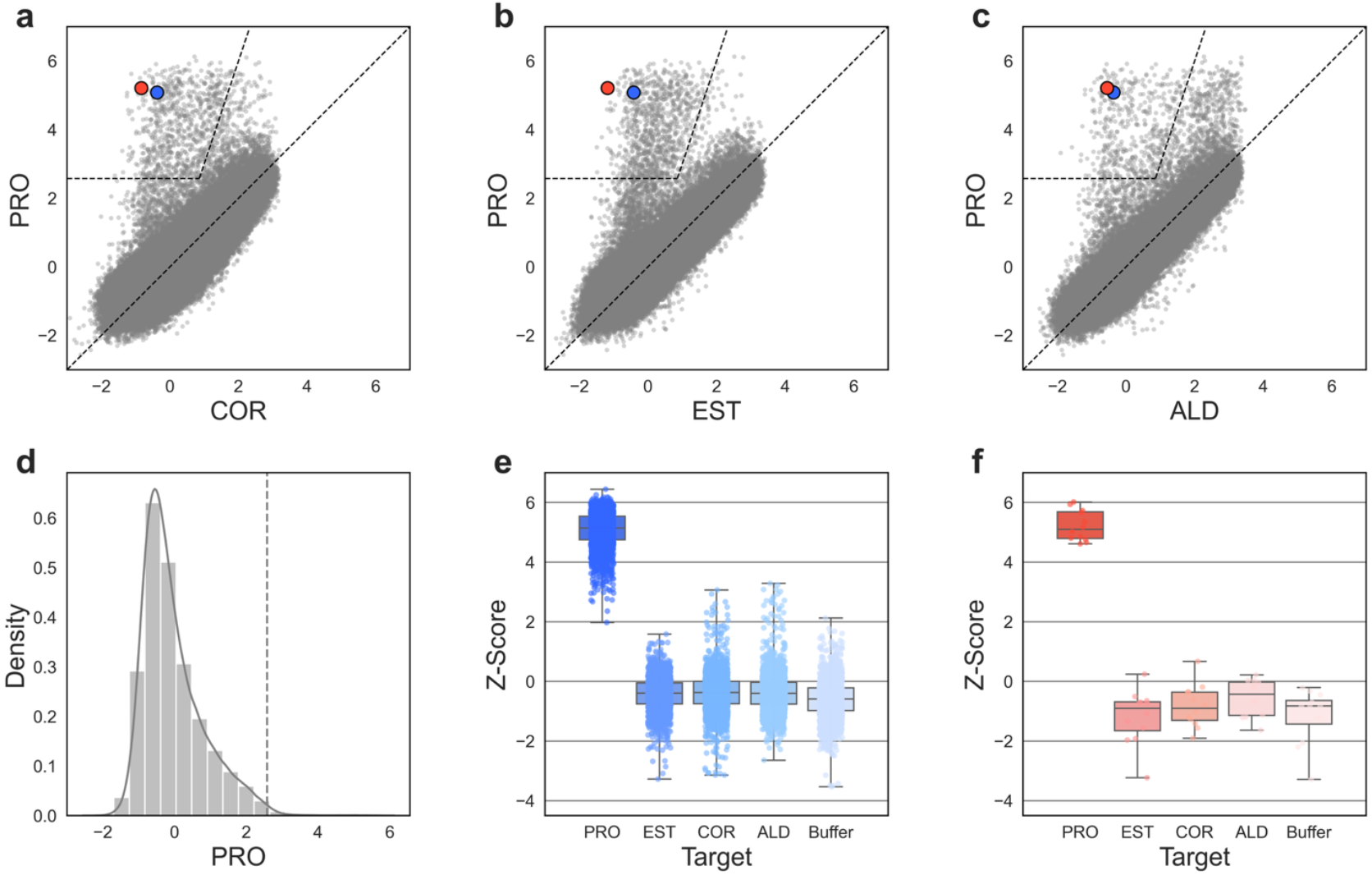
Specificity assessment of the aptamer pool for progesterone and three other steroids. (**a**–**c**) Z-score comparisons between target pairs for all sequences detected on the flow cell. Each point represents the average Z-score for a single sequence across one or more replicates. The areas at top left, demarcated by dashed lines, show sequences that are likely to bind to the target shown on the y-axis (Z-score ≥ 2.576) with high specificity (Z-score ratio ≥ 3). Z-scores for PRO-01 and PRO-02 are indicated as a blue dot and a red dot in each of the scatter plots. (**d**) Probability density function of Z-scores for progesterone, where the y-axis indicates the likelihood that any given sequence would yield a particular Z-score, as indicated along the x-axis. (**e**,**f**) Z-score distributions for PRO-01 (blue gradient) and PRO-02 (red gradient) in the presence of buffer or each of the four steroid analytes. Box plots show aptamer cluster-level data for PRO-01 (796 clusters) and PRO-02 (4 clusters). Medians are indicated by box centers; box edges represent quartiles; whiskers denote range.

By comparing Z-scores across analytes, we identified aptamers with strong responses to progesterone but minimal responses to analogs. We defined “high-affinity” candidates as those with Z-scores ≥ 2.576 for progesterone (**Fig. 2a–d**). To ensure specificity, we imposed a threshold of Z-score ≤ −0.3 for all non-target steroids. This stringent cutoff excludes sequences with modest cross-reactivity, reducing false positives from weak or non-specific binding. From the pool of sequences meeting both criteria, we selected the top two candidates (PRO-01 and PRO-02) based on their abundance on the flow cell and the magnitude of their on-target versus off-target Z-scores (**Table 1**). PRO-01 and PRO-02 achieved Z-scores of 5.1 and 5.2 for progesterone, while maintaining maximum off-target Z-scores of −0.3 and −0.5, respectively (**Fig. 2e,f**).

**Table 1.**
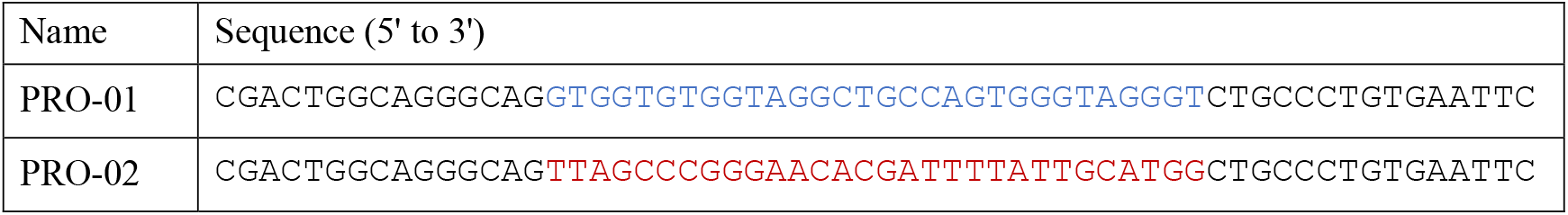
Sequences of high-specificity DNA aptamers for progesterone. Blue or red letters represent the randomized N(30) region; fixed forward and reverse primer regions flank both sides.

### Characterization of selected progesterone aptamers

To validate and characterize the binding performance of PRO-01 and PRO-02, we conducted further experiments in solution. We synthesized DNA aptamer strands containing the full 60-nt sequence used during selection, which includes a central 30-nt randomized region flanked by 15-nt forward and reverse primer regions (**Table S1**). A Cy3 fluorophore was attached to the 5’ end of the aptamer. These strands were then hybridized with a 3’ end quencher-labeled displacement strand complementary to the forward primer region, which resulted in approximately 90% fluorescence quenching upon annealing (**Fig. S1**). Target binding induces a conformational change that displaces the quencher strand, resulting in increased fluorescence. After generating binding curves against progesterone, we determined that PRO-01 and PRO-02 achieved *K*_*D*_ values of 28 nM and 36 nM (**Fig. 3a,b**). For a detailed description of the *K*_*D*_ calculation, see **Supporting Information, Section i.h**. This sensitivity covers progesterone’s physiologically relevant range, which fluctuates over short timeframes between low and high nanomolar concentrations in ovulatory women^16,17^. Notably, PRO-02 demonstrated superior specificity, with negligible responses to estradiol, aldosterone, and cortisol even at micromolar concentrations (**Fig. 3b**). Together, these results suggest that both aptamers are promising for progesterone sensing applications, with PRO-02 offering the added benefit of enhanced selectivity against structurally related steroid hormones.

**Figure 3.**
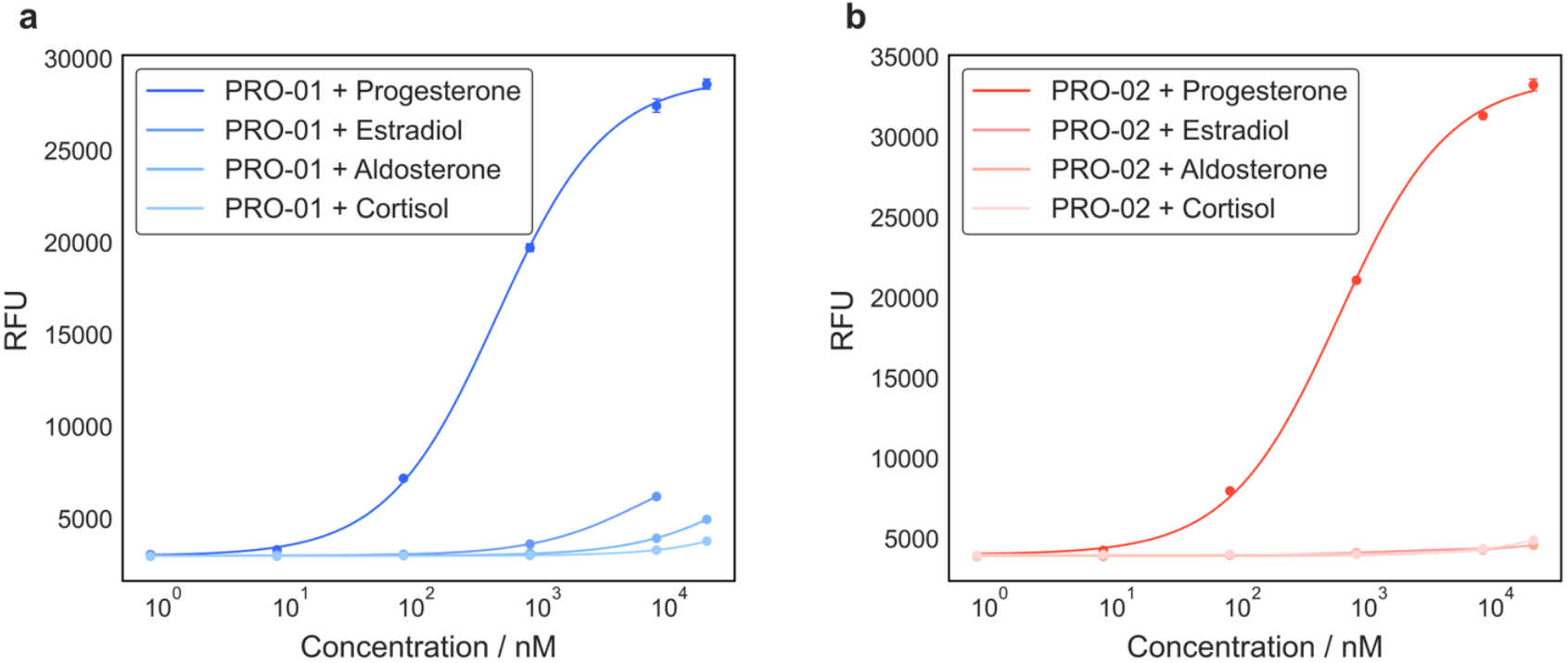
Characterization of our top-performing progesterone aptamer candidates, PRO-01 and PRO-02. Binding curves for (**a**) PRO-01 (blue gradient) and (**b**) PRO-02 (red gradient) with progesterone and steroid analogs in solution. Data points represent means from three separate experiments, with error bars showing the standard deviation. For estradiol, aptamer responses were assessed at only five concentrations due to solubility constraints that limited testing at higher target concentrations.

## Conclusion

In this study, we identified DNA aptamers with both high affinity and exceptional specificity for the steroid hormone progesterone. Using our high-throughput array platform, we were able to rapidly screen millions of aptamer sequences in parallel and assess their binding performance against multiple structurally similar analogs. PRO-01 achieved a *K*_*D*_ of 28 nM for progesterone, and PRO-02 achieved a *K*_*D*_ of 36 nM with negligible cross-reactivity to other major steroid hormones including cortisol, estradiol, and aldosterone.

These aptamers are fully synthetic and chemically stable, making them well-suited for downstream engineering into various biosensing formats, such as structure-switching aptamer beacons^22^. The same discovery pipeline can be readily extended to other small-molecule targets for which the development of high-specificity aptamers has been historically challenging, such as metabolic intermediates, drug isomers, or environmental contaminants. In particular, aptamers like PRO-02 may serve as core molecular recognition elements in diagnostic platforms for hormone monitoring, fertility tracking, and therapeutic drug response. Overall, the discovery of high-specificity DNA aptamers for progesterone using our integrated selection and screening strategy establishes a robust foundation for creating next-generation biosensors with clinical and research relevance.

## Acknowledgement

We thank Dr. Eric T. Kool (Stanford University) for valuable discussions. H.F. acknowledges support from Stanford Graduate Fellowship in Science and Engineering (SGF) and Funai Overseas Scholarship (FOS).

## Author contributions

Conceptualization: H.F., L.W., Y.G., H.T.S.

Methodology: H.F., L.W., Y.G., L.A.H.

Investigation: H.F., L.W., Y.G., L.A.H.

Visualization: H.F., L.W.

Funding acquisition: H.T.S.

Project administration: H.T.S.

Writing – original draft: H.F., H.T.S.

Writing – review & editing: all authors.

## Data and materials availability

All data are available in the main text or the supplementary materials.

