## Supporting Information for "Discovery of high-specificity DNA aptamers for progesterone using a high-throughput array platform"

**Table of Contents**

i. **Materials and Methods**

i.a **Materials.**

i.b **Overview of sequence design.**

i.c **Preparation of buffer and target solutions.**

i.d **Selection of progesterone aptamer using capture-SELEX.**

i.e **Preparation for high-throughput sequencing and screening.**

i.f **High-throughput sequencing and specificity screen.**

i.g **Processing data from specificity screen.**

i.h **Characterization of aptamers via plate reader.**

ii. **References**

### i. Materials and Methods

#### i.a Materials.

1 M Tris-HCl buffer (pH 7.5) (Cat# BP1757), GelStar (Cat# BMA50535), phosphate-buffered saline (Cat# BP399), 1 N NaOH (Cat# SS266-1), 1 M $MgCl_2$ (Cat# AM9530G), streptavidin agarose resin (Cat# 20353), and DNA Gel Loading Dye (6X) (Cat# R0611) were purchased from Thermo Fisher Scientific. 5 M NaCl (Cat# S5150), anhydrous dimethyl sulfoxide (DMSO) (Cat# 276855), Tween 20 (Cat# P9416), 3 M NaOAc (Cat# 567422), progesterone (Cat# P0130), estradiol (Cat# E2758), aldosterone (Cat# A9477), cortisol (hydrocortisone, Cat# H0888), and cholesterol (Cat# C8667) were purchased from Sigma-Aldrich. 1M KCl (Cat# P035) and 1M $CaCl_2$ (Cat# C0477) were purchased from Teknova. HyPure molecular biology-grade water (Cat# SH31191) was acquired from Cytiva. Novex Tris-borate-EDTA (TBE) Gels, 10% (EC6275) were purchased from Invitrogen. 20-bp molecular ruler (Cat# 1708201) was purchased from Bio-Rad. All reagents for DNA sequencing including Nextera XT DNA Library Preparation Kit were obtained from Illumina. Axygen AxyPrep Mag PCR Clean-up Kit (Cat# 14-223-227) and 96-well half-area black microplate (Cat# 3694) were purchased from Corning. The MinElute PCR Purification Kit (Cat# 28004) was purchased from QIAGEN. GoTaq G2 Hot Start Master Mix (Cat# M7433) was purchased from Promega. 0.2-µm syringe filters were purchased from VWR. 10-kDa molecular-weight cutoff size-exclusion columns were purchased from MilliporeSigma. EcoRI (high fidelity, Cat# R3101) and associated buffers were obtained from New England Biolabs. Oligonucleotides shown in **Table S1** were purchased from Integrated DNA Technologies.

#### i.b Overview of sequence design.

The DNA sequences used in this study were designed to support both the capture-SELEX process and high-throughput screening on a modified Illumina MiSeq platform (**Table S1**). Each oligonucleotide consisted of a central 30-nucleotide (nt) randomized region flanked by 15-nt forward and reverse primer regions. During SELEX, the forward primer was labeled with 5(6)-carboxyfluorescein (FAM) to enable monitoring of enrichment, and the reverse primer was biotinylated to facilitate strand separation via streptavidin beads. For high-throughput sequencing, Illumina adaptor and index sequences were appended to both ends of the aptamer construct. As a result, each DNA molecule immobilized on the MiSeq flow cell contains the following elements in order (5′ to 3′): index, forward adaptor, forward primer, randomized region, reverse primer, reverse adaptor, and second index. Only the core region (forward primer, randomized region, and reverse primer) was used during SELEX and is relevant for binding affinity measurements. The reverse adaptor and index sequences were removed using EcoRI digestion before target-binding analysis. A complete list of all oligonucleotide sequences and modifications, including primers, adaptors, capture strands, and displacement strands, is provided in **Table S1**.

**Table S1.** DNA sequences used in this study.

| Category | Name | Sequence (5' to 3') |
|---|---|---|
| Capture SELEX | Library | CGACTGGCAGGGCAG-N(30)-CTGCCCTGTGAATTC |
| | Biotinylated reverse primer | Biotin-GAATTCACAGGGCAG |
| | FAM-labeled forward primer | FAM-CGACTGGCAGGGCAG |
| | Biotinylated capture strand | CTGCCCTGCCAGTCG-(18-atom hexa-ethyleneglycol linker)-biotin |
| High-throughput sequencing and screening | Sequencing adaptor for forward primer | TCGTCGGCAGCGTCAGATGTGTATAAGAGACAGNNNNCGACTGGCAGGGCAG |
| | Sequencing adaptor for reverse primer | GTCTCGTGGGCTCGGAGATGTGTATAAGAGACAGGAATTCACAGGGCAG |
| | 15-mer Cy3-labeled displacement strand | CTGCCCTGCCAGTCG-Cy3 |
| | EcoRI complementary strand | AGAGACAGGAATTC |
| Plate reader assay | Aptamer labeled with Cy3 at the 5' end | Cy3-CGACTGGCAGGGCAG-N(30)-CTGCCCTGTGAATTC |
| | 13-mer displacement strand labeled with BHQ2 at the 3' end | CTGCCCTGCCAGT-BHQ2 |

#### i.c Preparation of buffer and target solutions.

Stock solutions were prepared as follows: 100 mM cortisol in DMSO, 50 mM progesterone in DMSO, 20 mM estradiol in DMSO, 50 mM aldosterone in DMSO, and 1 mM cholesterol in ethanol. Stock solutions were stored tightly sealed at 4°C for short-term use (within several weeks) or at -20 °C for long-term storage.

#### i.d Selection of progesterone aptamer using capture-SELEX.

Our capture-SELEX was based on an existing protocol[1,2]. Each round of selection used a five-fold molar excess of biotinylated capture strand relative to the aptamer DNA in 250 µL of selection buffer containing 20 mM Tris-HCl, 120 mM NaCl, 5 mM KCl, 1 mM $MgCl_2$, 1 mM $CaCl_2$, and 0.01% Tween-20 in nuclease-free water. We used 1 nmol of the DNA library in the first round, and 100–500 pmol of single-stranded DNA in subsequent rounds. To initiate hybridization, the aptamer pool was heated together with the biotinylated capture strand at 95 °C for 5 minutes, then cooled to room temperature for at least 30 minutes to allow folding. The selection process began with loading 250 µL of streptavidin resin into a column, followed by five washes with selection buffer, discarding the flow-through after each wash. Unless otherwise specified, all washes were performed using 250 µL volumes. The annealed aptamer–capture strand library was passed through the column, and the resulting flow-through was passed through the same column three additional times to maximize capture efficiency. After capturing the library, the column was washed with selection buffer to remove unbound or weakly bound strands. To increase selection stringency, the number of washes was increased from 10 in rounds 1–3 to 12 in rounds 4–7.

We prepared a mixed steroid solution by combining stock solutions to yield final concentrations of 50 µM progesterone, 10 µM estradiol, 100 µM aldosterone, 100 µM cortisol, and 200 nM cholesterol in 750 µL of selection buffer. This solution was added to the column in three separate 250 µL portions. All steroid targets were introduced together in every round of SELEX. Although we performed subsequent specificity screens for each steroid, we ultimately excluded cholesterol from downstream analysis due to its comparatively low concentration. Target solutions were prepared by diluting the concentrated steroid stocks in ethanol or DMSO into selection buffer. Target concentrations were kept constant across all SELEX rounds. Starting from the fourth round, we collected the flow-through and measured fluorescence using a Qubit Fluorometer (Thermo Fisher Scientific) to monitor aptamer enrichment. Fluorescence was measured using a blue excitation filter (λ: 430–495 nm) and a green emission filter (λ: 510–580 nm) of the Qubit Fluorometer.

To determine the optimal number of PCR cycles for each round, we performed pilot PCR reactions. The PCR mixture consisted of 20 µL of 100 µM FAM-labeled forward primer, 20 µL of 100 µM biotinylated reverse primer, 1 mL of GoTaq Master Mix, and nuclease-free water to a final volume of 2 mL. This mixture was aliquoted into twenty 100 µL PCR tubes. PCR was performed using an Eppendorf Mastercycler X50 with the following thermal cycling conditions: initial denaturation at 95 °C for 2 minutes, followed by nine cycles of 95 °C for 15 seconds, 54 °C for 15 seconds, and 72 °C for 30 seconds, followed by 72˚C for 1 minute and then holding at 4˚C. All tubes were first subjected to four PCR cycles, after which 4 µL was taken from each tube and pooled (total volume: 80 µL). This pooled sample underwent four additional cycles. After each 4-cycle increment, 4 µL aliquots were collected for analysis, continuing up to 24 total cycles. PCR products from various cycle numbers (e.g., 4, 8, 12, 16, 20, 24 cycles) were analyzed by electrophoresis on a 10% TBE gel, with a 20-bp DNA ladder loaded at both ends for size reference. We monitored the appearance of a distinct product band at ~60 bp, corresponding to the combined length of fixed primer regions and the randomized region. Smearing at higher cycle numbers indicated overamplification. To balance yield and specificity, we selected the highest cycle number that gave a sharp product band with minimal smearing. The optimal cycle numbers varied by round: 12 (round 1), 11 (rounds 2–3), 10 (round 4), 13 (round 5), and 10 (rounds 6–7).

Following PCR amplification, the pooled reaction were purified using the MinElute PCR Purification Kit. Five volumes of Buffer PB (9.75 mL) and ~100 µL of 3 M NaOAc were added to adjust the pH (indicated by a yellow color change), and the mixture was thoroughly combined. The total volume was divided across four MinElute columns and centrifuged at 17,900 × g for 1 minute at 4 °C, repeating until the entire sample passed through. Columns were then spun again to remove residual liquid. Each column was washed with 750 µL Buffer PE, spun for 1 minute, and then dry-spun again. DNA was eluted in 20 µL of nuclease-free water per column, incubated for 1 minute before spinning. This elution was repeated two additional times, and eluates were pooled (~60 µL per column).

To convert the PCR amplicons into single-stranded DNA (ssDNA), 300 µL of Dynabeads™ MyOne™ Streptavidin C1 (SA-C1) beads were used. The beads were first resuspended and rotated for at least 10 minutes at room temperature to ensure a uniform suspension. They were then pulled down using a magnetic rack and allowed to settle for 1 minute. The beads were washed three times with 400 µL of 20 mM NaOH, each followed by a 15-minute incubation, to remove residual biotinylated oligos and contaminants. After the final NaOH wash, the beads were washed three times with 800 µL of selection buffer, mixing gently by hand rotation to equilibrate the beads. Next, the purified double-stranded DNA (dsDNA) library, containing a biotinylated reverse primer and a FAM-labeled forward primer, was added to the beads in selection buffer to a final volume of 700 µL. The bead–library mixture was incubated at room temperature for 1 hour to allow efficient binding between the biotinylated strand and the streptavidin-coated beads. Following binding, the beads were washed twice with 800 µL of selection buffer. The first supernatant was retained for quality control or troubleshooting. An additional wash was performed with 800 µL of nuclease-free water to remove residual salts and buffer components. To elute the unbiotinylated strand and recover ssDNA, 200 µL of 20 mM NaOH was added to the beads and the mixture was incubated for at least 8 minutes at room temperature. The beads were then pulled down with a magnetic rack, and the supernatant containing the released ssDNA was transferred to a new 1.5 mL tube. To neutralize the solution, 35 µL of 1 M Tris-HCl (pH 7.0) was added, followed by 500 µL of selection buffer. The ssDNA was then purified and concentrated using centrifugal filters with a 10 kDa molecular-weight cutoff. Samples were centrifuged at 14,000 × g for 20 minutes. Buffer exchange was performed twice using 400 µL of nuclease-free water, spinning at 14,000 × g for 20 minutes each time. Finally, the column was flipped into a clean 1.5 mL tube and centrifuged at 1,000 × g for 1 minute to recover the concentrated ssDNA. DNA concentration was quantified using both a NanoDrop spectrophotometer and a Qubit Fluorometer (ssDNA assay kit).

**i.e Preparation for high-throughput sequencing and screening.**

Single-stranded DNA was added to a final volume of 300 µL PCR mix, which also included distinct sequencing adaptors for the forward and reverse primers. This mixture was amplified using the same PCR protocol described for capture-SELEX. After four PCR cycles, the adaptor-ligated products were purified using the Axygen AxyPrep Mag PCR Clean-up Kit. Next, the ssDNA was added to a final volume of 450 µL PCR mix and indexed using the Nextera XT DNA Library Preparation Kit. The indexed pool was again cleaned using the AxyPrep Mag PCR Clean-up Kit. The cleaned DNA was then run on a 10% TBE gel. Desired DNA bands were identified, excised, and crushed. The gel fragments were incubated overnight in 400 µL of Tris-EDTA (TE) buffer at room temperature. The supernatant was separated from the gel pieces by centrifugation through 0.2 µm VWR syringe filters at 14,000 × g for 3 minutes. The filtered DNA was concentrated using 10 kDa size-exclusion columns, and the buffer was exchanged to water. The column was flipped into a clean 1.5 mL tube and spun at 1,000 × g for 1 minute to recover the concentrated DNA. DNA concentration was determined using both a NanoDrop spectrophotometer and a Qubit Fluorometer.

#### i.f High-throughput sequencing and specificity screen.

To perform high-throughput screening of aptamers, we used a modified Illumina MiSeq system (previously described in Refs. 2,3) to screen aptamer clusters. The MiSeq performs bridge amplification of the aptamer pool to generate monoclonal DNA clusters, each comprising approximately 1,000 identical strands. During the first read, the MiSeq determines both the sequence and spatial coordinates of each cluster on the flow cell. Instead of proceeding to the second read, we repurposed these cycles to deliver custom reagents, including 15-mer Cy3-labeled displacement strand and steroid targets. Each DNA construct on the flow cell includes, in 5′ to 3′ order: the first index, forward adaptor, forward primer, randomized region, reverse primer, reverse adaptor, and second index. For affinity measurements, only the core aptamer sequence (forward primer, randomized region, and reverse primer) is functionally relevant, as this is the portion present during SELEX. The reverse adaptor and second index were removed by EcoRI digestion with an EcoRI complementary strand prior to target binding analysis.

The screening process consisted of alternating buffer and target exposure cycles to monitor displacement of the labeled strand upon target binding. During each buffer cycle, residual strands and ligands were removed using 0.05 M NaOH with 0.25% SDS, followed by a wash with selection buffer. Next, 200 nM 15-mer Cy3-labeled displacement strand in selection buffer was introduced, and the flow cell was heated to 80 °C and gradually cooled to 22 °C over ~30 minutes to promote hybridization. A final wash with selection buffer was performed before imaging. In each target cycle, the flow cell was first washed with selection buffer, then exposed to increasing concentrations of the target solution prepared in selection buffer. Target concentrations matched those used in the capture-SELEX process: 50 µM progesterone, 10 µM estradiol, 100 µM aldosterone, 100 µM cortisol, and 200 nM cholesterol. All buffer and target measurements were conducted in triplicate.

#### i.g Processing data from specificity screen.

To assess specificity, we calculated Z-scores that quantify the relative change in fluorescence upon target binding[2]. For each measurement cycle, the percent change in fluorescence intensity for each aptamer cluster was calculated as:

$$\%_{change} = \frac{I_{buffer} - I_{target}}{I_{buffer}} \qquad \text{(eq. S1)}$$

We then normalized these percent changes across clusters within a cycle to account for variability due to cluster degradation or signal drift over time. Z-scores were computed using the formula:

$$Z = \frac{\%_{change,\ cluster} - \mu_{cycle}}{\sigma_{cycle}} \qquad \text{(eq. S2)}$$

where $\mu_{\text{cycle}}$ and $\sigma_{\text{cycle}}$ represent the mean and standard deviation of the percent changes for all clusters within that cycle. These Z-scores were then averaged across replicates for each aptamer–target pair to yield a single Z-score per sequence per target. To define target-binding aptamers, we applied a significance threshold to the Z-score for the target, denoted as $z_t$, requiring $z_t > 2.576$. Specificity was further assessed at the ratio of the Z-score for the target relative to that for the non-target molecule. Only aptamers that satisfied both $z_{target} > 2.576$ and $\frac{z_{target}}{z_{off-target\ max}} > 3$ — where $z_{off-target\ max}$ is the maximum Z-score across all off-targets and buffer — were considered target-specific. This two-tiered filtering process ensured high-confidence identification of monospecific sequences suitable for downstream analysis. We also applied a filter to exclude fluorescent signals below 750 and above 1500 during the final displacement strand cycle of the high-throughput aptamer array screening as these values are indicative of instrument failure to accurately measure fluorescence. Additionally, we removed populations showing a percent change in fluorescence intensity >0.5 or <-0.3 during the buffer cycle, as these sequences consistently responded across all target cycles irrespective of conditions. We excluded populations showing >0.5 percent change during any off-target cycle or <-0.3 across all target cycles.

#### i.h Characterization of aptamers via plate reader

We used a previously established two-step binding model to determine the overall dissociation constant ($K_D$) of each aptamer for its target molecule[1,4,5]. In this model, $K_D$ is computed as the ratio of two independently measured constants: $K_D = K_{D,1}/K_{D,2}$ where $K_{D,1}$ represents the dissociation constant for hybridization between the aptamer and the displacement strand (*i.e.,* aptamer + displacement strand ⇌ aptamer · displacement strand), and $K_{D,2}$ represents the dissociation constant for target binding to the aptamer–displacement strand complex, which induces strand displacement and fluorescence recovery (*i.e.,* aptamer·displacement strand + target ⇌ aptamer·target + displacement strand). To determine $K_{D,1}$, we generated a binding curve by incubating 50 nM aptamer labeled with Cy3 at the 5' end with increasing concentrations of the 13-mer displacement strand labeled with BHQ2 at the 3' end (**Fig. S1**). The displacement strand concentration that achieved ~90% fluorescence reduction was selected for the second assay. Based on these binding curves, we selected final displacement strand concentrations of 2500 nM for PRO-01 and 500 nM for PRO-02. These concentrations were then used in the second binding assay to determine $K_{D,2}$. Specifically, 50 nM Cy3-labeled aptamer was pre-annealed with the designated displacement strand concentration, followed by incubation with increasing concentrations of the target steroid for 30 minutes (**Fig. 3**). Fluorescence recovery was measured using a black half-area 96-well plate and a microplate reader (Synergy H1, BioTeK) equipped with a filter cube (excitation: 538/63; emission: 590/35; gain: 55). All measurements were conducted in triplicate at room temperature in selection buffer. Data were analyzed in Python to fit to the Hill equation (with a fixed Hill coefficient of $n = 1$) using the "curve_fit" function from the SciPy library.

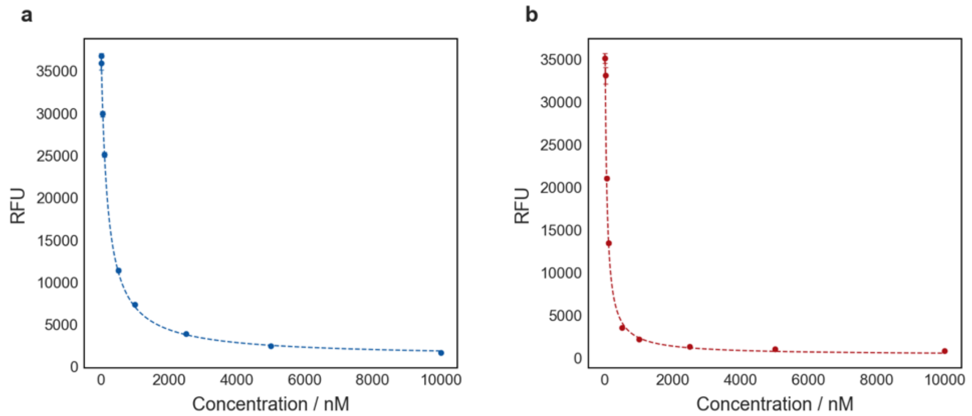


**Figure S1.** Binding curves showing fluorescence quenching of 50 nM aptamers labeled with Cy3 at the 5' end for (**a**) PRO-01 and (**b**) PRO-02 upon incubation with increasing concentrations of the 13-mer displacement strand labeled with BHQ2 at the 3' end.
